# Assessment of ethanol-induced toxicity on iPSC-derived human dopaminergic neurons using a novel high-throughput mitochondrial neuronal health (MNH) assay

**DOI:** 10.1101/2020.08.12.237461

**Authors:** Annika Zink, Josefin Conrad, Narasimha Swami Telugu, Sebastian Diecke, Andreas Heinz, Erich Wanker, Josef Priller, Alessandro Prigione

## Abstract

Excessive ethanol exposure can cause mitochondrial and cellular toxicity. In order to discover potential counteracting interventions, it is essential to develop assays capable of capturing the consequences of ethanol exposure in human dopaminergic (DA) neurons, which are crucial for the development and maintenance of alcohol use disorders (AUD). Here, we developed a novel high-throughput (HT) assay to quantify mitochondrial and neuronal toxicity in human DA neurons from induced pluripotent stem cells (iPSCs). The assay, dubbed <u>m</u>itochondrial <u>n</u>euronal <u>h</u>ealth (MNH) assay, combines live-cell measurement of mitochondrial membrane potential (MMP) with quantification of neuronal branching complexity post-fixation. Using the MNH assay, we demonstrated that chronic ethanol exposure in human iPSC-derived DA neurons decreases MMP and branching complexity in a dose-dependent manner. The toxic effect of ethanol on DA neurons was already detectable after 1 hour of exposure, and occurred similarly in DA neurons derived from healthy individuals and from patients with AUD. We next used the MNH assay to carry out a proof-of-concept compound screening using FDA-approved drugs. We identified potential candidate drugs modulating acute ethanol toxicity in human DA neurons. Among these drugs, flavoxate and disulfiram influenced mitochondrial neuronal health independently from ethanol, leading to amelioration and worsening, respectively. Altogether, we developed an HT assay to probe human mitochondrial neuronal health and used it to assess ethanol neurotoxicity and to identify modulating agents. The MNH assay represents an effective new tool for discovering modulators of mitochondrial neuronal health and toxicity in live human neurons.

## Introduction

Ethanol is the most frequently abused drug worldwide and its excessive consumption is a leading risk factor for disability and death (1). High ethanol consumption over time can lead to serious health and social problems, including the development of alcohol use disorder (AUD). AUD is among the most prevalent mental disorders in industrialized countries (2). Once ingested, ethanol quickly distributes throughout the body and reaches the brain within minutes. Given this rapid and vast spread, ethanol can cause direct organ toxicity. Ethanol-induced neurotoxicity is particularly detrimental given that damaged neurons cannot be replaced. Within the central nervous system, ethanol exposure directly affects dopaminergic (DA) neurons, resulting in increased extracellular dopamine mainly in the striatum (3) higher firing frequency and increased excitability (4), as well as decreased DA synthesis and dopamine D2 receptor availability in AUD patients (5).

At the cellular level, an important role during ethanol intoxication may be played by mitochondria, crucial organelles that safeguard cellular homeostasis. Mitochondria provide energy in form of ATP through oxidative phosphorylation (OxPhos), which turns ADP into ATP by releasing the energy stored in the proton gradient also known as mitochondrial membrane potential (MMP) (6). Mitochondria are also crucial for cell death (7). Selective MMP permeabilization activates caspase-driven apoptotic cell death through opening of the mitochondrial permeability transition pore (mPTP) (8). MMP is thus an essential parameter for viable cells, since it is important for both ATP generation and initiation of apoptosis, and may serve as a marker of cell health (9). Various studies reported ethanol-induced toxicity in mitochondria located in the brain (10), including increased mitochondrial production of free radicals (11), alterations in mitochondrial respiration and MMP (11,12), impairment of ATP production (13,14), and cell death induction (13,15).

The mechanisms underlying ethanol-induced toxicity in human brain cells remain largely unknown. Most investigations are based on animal models, which may not fully recapitulate the human disease state, on post-mortem tissues, which correlates more to an endstage of AUD, or on patient brain imaging, which provides macroscopic data lacking information at the cellular and molecular level (16). Given the lack of suitable human cell-based model systems for the assessment of neurotoxicity and drug discovery, our knowledge of compounds capable of counteracting ethanol toxicity in humans is limited.

Here, we used human iPSCs to investigate the toxic consequences of ethanol in human DA neurons. In order to assess human neurotoxicity in a quantitative and high-throughput (HT) manner, we devised a new assay that we named <u>m</u>itochondrial <u>n</u>euronal <u>h</u>ealth (MNH) assay. The MNH assay is based on high-content imaging (HCI) and combines live-cell measurement of MMP with quantification of neuronal branching outgrowth. Using this assay, we demonstrated the acute and chronic effects of ethanol toxicity on mitochondrial neuronal health in human DA neurons that we derived from control individuals and from subjects with AUD. We also showed that the MNH assay can be used to perform compound screenings to identify drugs influencing mitochondrial neuronal health. Hence, the newly developed MNH assay represents an effective HT tool for analyzing the cellular health of human iPSC-derived neurons and for discovering potential modulatory interventions.

## Results

### Development of the MNH assay

In order to assess the toxic effects of ethanol in human neurons, we first generated neural progenitor cells (NPCs) from healthy control-derived iPSCs (XM001 line) (17) and human embryonic stem cells (hESCs) (H1 line) (18) using a small molecule-based protocol (19). We then differentiated NPCs into post-mitotic neurons enriched for DA neurons **(Fig. 1A)**. The pluripotent stem cell (PSC)-derived DA neurons expressed the neuron-specific marker MAP2 and DA markers tyrosine hydroxylase (TH) and FOXA2 **(Fig 1B)**. In the differentiated cultures, 75% of cells expressed the pan-neuronal marker beta tubulin-III (TUJ1) and 20% expressed TH **(Fig. 1C)**. These numbers are in accordance with previous protocols (20). We monitored neuronal network activity using micro-electrode array (MEA). We confirmed that the generated DA neurons were functional and exhibited multiple spontaneous spikes after 4-8 weeks in culture **(Fig. 1D)**.

**Fig. 1.**
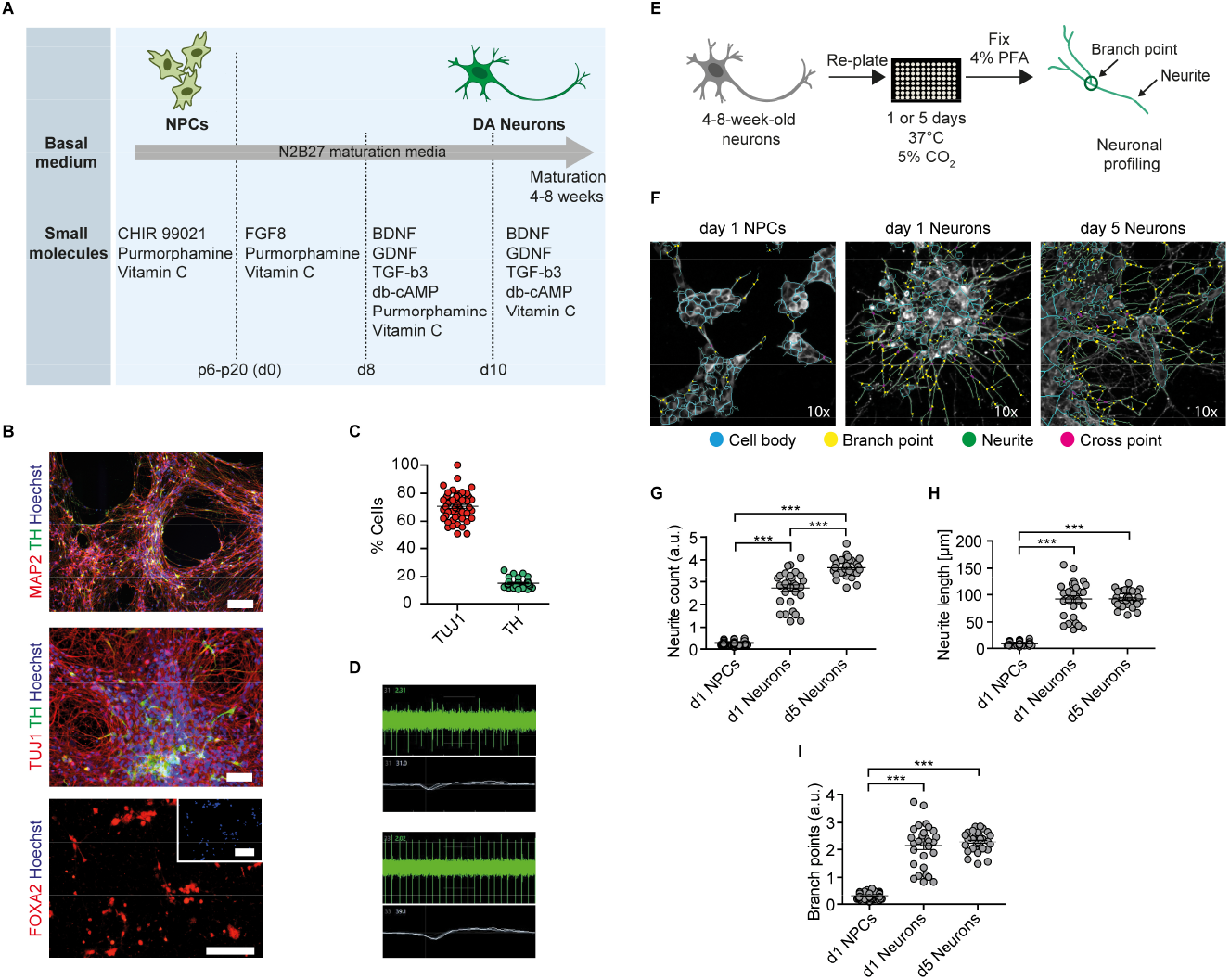
Generation of functional DA neurons and HCI-based neuronal profiling. **(A)** Schematics of the iPSC-based generation of NPCs and DA neurons, indicating the timepoints of different media formulations. **(B)** 4-8-week-old DA neurons generated from control iPSCs (XM001 line) expressed the neuronal marker TUJ1 and MAP2, and the DA-specific markers FOXA2, and TH. Scale bar: 100 μm. **(C)** HCI-based quantification of control DA neurons (XM001) expressing TUJ1 and TH at 8 weeks of culture (mean +/- SEM). Each dot represents mean values (% neuronal cells) from one well. **(D)** Representative recordings (spike plot and wave plot) of network-based electrophysiological properties of control DA neurons after 8 weeks in culture using MEA. **(E)** Schematic workflow of HCI-based neuronal profiling analysis. **(F)** Representative images of NPCs and DA neurons (at 4-8 weeks of differentiation) fixed after day 1, and of DA neurons fixed after day 5. Cell bodies depicted in blue, branch points in yellow, neurites in green, and cross points in magenta. All images were taken at 10x magnification. **(G-I)** Quantification of neurite count **(G)**, neurite length **(H)**, and branch points **(I)** in NPCs and 4-8-week-old DA neurons at day 1 and day 5 after re-plating (30 technical replicates for DA neurons and 60 technical replicates for NPCs; each dot represents the value obtained from one well; mean +/- SEM; ***p<0.001; one-way ANOVA followed by Dunnett’s multiple comparison test).

We next established an HCI-based approach to assess mitochondrial function and neuronal growth capacity in DA neurons. We dubbed the method <u>m</u>itochondrial <u>n</u>euronal <u>h</u>ealth (MNH) assay. To assess neuronal branching capacity, we cultivated DA neurons for 4-8 weeks, re-plated them on the assay plate and kept them for 1 day or 5 days after plating. We then fixed the cultures and stained them with TUJ1 to visualize neuronal arborization **(Fig. 1E)**. Since TUJ1 staining does not allow to discriminate between axons and dendrites, we chose to refer to any projection from the cell body as neurite. In order to assess differences in neurite outgrowth, we compared DA neurons fixed 1 day or 5 days post-plating to NPCs fixed 1day post-plating **(Fig. 1F)**. The MNH assay effectively captured and quantified differences in neuronal arborization. DA neurons at days 5 post-plating showed increased neurite count **(Fig. 1G)**, neurite length **(Fig. 1H)**, and branch points **(Fig. 1I)** compared to neurons at day 1 post-plating or NPCs at day 1 post-plating. The cellular population appeared also more homogenous in day 5 DA neurons, as indicated by lower variability in the distribution of the replicates **(Fig. 1G-I)**.

In order to assess MMP, we implemented an HCI assay that we previously established for iPSC-derived NPCs (21). We measured MMP using the potentiometric fluorescent dye tetramethylrhodamine methyl ester (TMRM), a lipophilic cation that accumulates in the mitochondrial matrix in proportion to the potential of the membrane. We re-plated 4-8-week-old DA neurons on the assay plate and kept them for 5 days before live-staining for MMP and subsequent neuronal branching quantification **(Fig. 2A)**. Stimulation with the oxidative phosphorylation uncoupler FCCP and the complex III inhibitor Antimycin A (Ant.A) provoked a decrease in the MMP in a dose-dependent manner **(Fig. 2B)**, whereas the ATPase inhibitor Oligomycin led to a dose-dependent increase in the MMP **(Fig. 2C)**. The MMP-modulating effects of the mitochondrial inhibitors are in line with previous works (22,23). The use of stimulating and inhibiting MMP modulators enabled us to determine the z-factor, a statistical indicator of HT bioassays that is considered excellent if 0.5>x>1 (Zhang et al., 1999). The MNH assay had a z-factor of 0.747, suggesting excellent features regarding reproducibility and robustness **(Fig. 2C)**. Branching complexity did not change upon exposure to FCCP and Ant.A at a concentration of up to 100 μM **(Fig. 2D-F)**, or to Oligomycin at a concentration of up to 200 μM **(Fig. 2G-I)**. Taken together, the MNH assay was able to quantitatively identify changes in neuronal outgrowth capacity and in MMP-specific function.

**Fig. 2.**
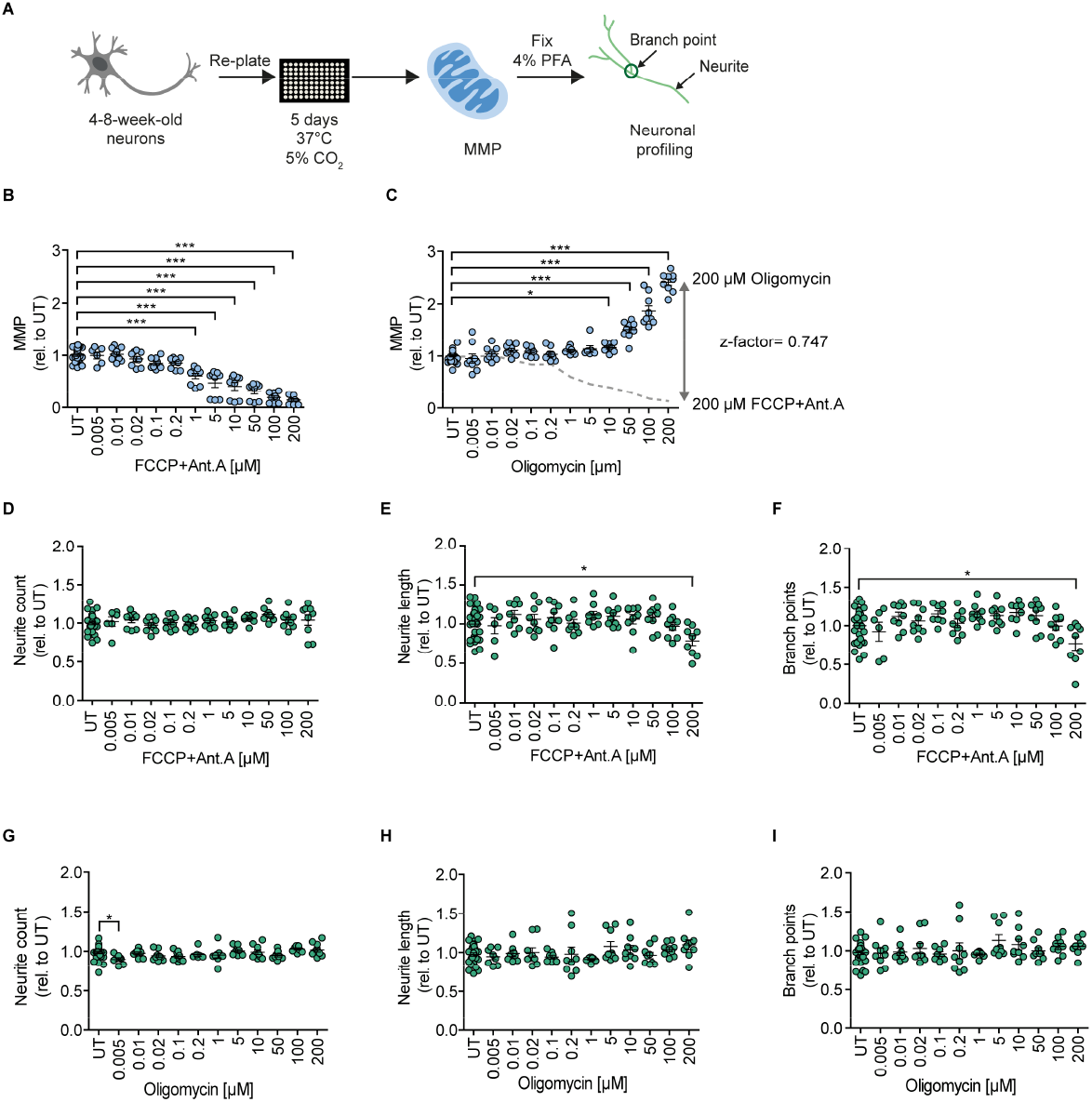
MNH assay development. **(A)** Schematics of the MNH assay. **(B)** Quantification of MMP in 4-8-week-old DA neurons derived from hESCs (H1 line) treated with increasing concentrations of FCCP and Ant. A (0.005 μM-200 μM). Each dot represents the average of the values obtained from one well normalized to the corresponding untreated (UT) control exposed to the assay media only (n= 3 independent experiments; mean +/- SEM; ***p<0.001; one-way ANOVA followed by Dunnett’s multiple comparison test). **(C)** Quantification of MMP in 4-8-week-old DA neurons from control hESCs (H1 line) treated with increasing concentrations of Oligomycin (0.005 μM-200 μM; mean +/- SEM; *p<0.05, ***p<0.001; one-way ANOVA followed by Dunnett’s multiple comparison test). Each dot represents the average of values obtained from one well normalized to the corresponding UT control exposed to the assay media only (n= 3 independent experiments). Assessment of the z-factor of 0.747 obtained at 200 μM FCCP and 200 μM Ant.A versus 200 μM Oligomycin suggesting an excellent assay. **(D-F)** Quantification of neurite count **(D)**, neurite length **(E)**, and branch points **(F)** in 4-8-week-old DA neurons from hESCs (H1 line) treated with increasing concentrations of FCCP and Ant.A (0.005 μM-200 μM; mean +/- SEM; *p<0.05; one-way ANOVA followed by Dunnett’s multiple comparison test). Each dot represents values obtained from one well (n= 3 independent experiments), normalized to the corresponding UT controls exposed to the assay media only. **(G-I)** Quantification of neurite count **(G)**, neurite length **(H)**, and branch points **(I)** in 4-8-week-old DA neurons from hESCs (H1) treated with increasing concentrations of Oligomycin (0.005 μM-200 μM; mean +/- SEM; *p< 0.05; one-way ANOVA followed by Dunnett’s multiple comparison test). Each dot represents values obtained from one well (n= 3 independent experiments), normalized to the corresponding UT controls exposed to the assay media only.

### The MNH assay detects chronic and acute ethanol-induced neurotoxicity in human DA neurons

In alcohol-naïve individuals blood alcohol concentrations (BACs) of 10 – 50 mM typically lead to sedation, motor incoordination and cognitive impairment. BACs of ≥ 100 mM cause strong sedation and can lead to coma or death in alcohol-naïve individuals. Instead, in chronic alcohol consumers BACs of up to 300 mM have been reported (24), and 100 – 200 mM typically lead to sedation, anxiolysis and hypnosis (25,26).

To address the toxic effects of ethanol on human DA neurons, we tested healthy DA neurons after 4-8 weeks of differentiation from iPSCs (XM001 line) and hESCs (H1 line). We exposed healthy DA neurons to different concentrations of ethanol in the media for 7 days (chronic exposure), with full media exchange every other day **(Fig. 3A)**. Chronic exposure to ethanol over 7 days increased MMP at ethanol concentrations of 10 mM-100 mM and caused the MMP to collapse at ethanol concentrations ≥ 500 mM **(Fig. 3B)**. Ethanol concentrations higher than 250 mM also strongly impaired neurite outgrowth **(Fig. 3C-E)**.

**Fig. 3.**
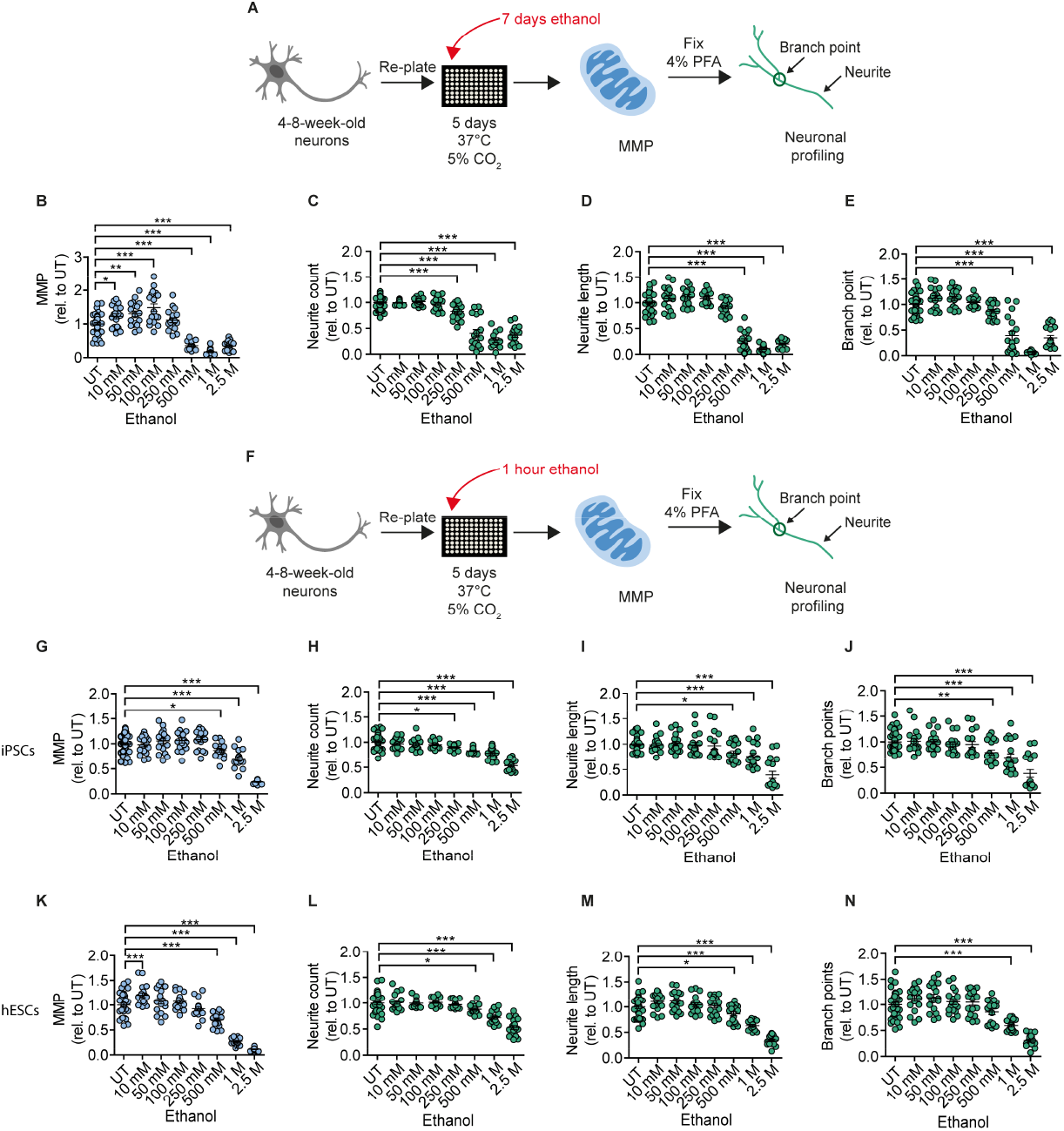
Chronic and acute ethanol exposure leads to decreased MMP and reduced branching complexity in iPSC- and hESC-derived DA neurons. **(A)** Schematic of MNH assay workflow for chronic (7 days) ethanol exposure. **(B)** Quantification of the MMP in 4-8-week-old DA neurons from control iPSCs (XM001) treated for 7 days (chronic) with increasing concentrations of ethanol (10 mM-2.5 M; mean +/- SEM; *p< 0.05, **p<0.01, ***p< 0.001; one-way ANOVA followed by Dunnett’s multiple comparison test). Each dot represents values obtained from one well (n= 3 independent experiments), normalized to the corresponding UT controls exposed to the assay media only. **(C-E)** Quantification of the neuronal branching complexity including neurite count **(C)**, neurite length **(D)**, and branch points **(E)** in 4-8-week-old DA neurons from control iPSCs (XM001) treated with increasing concentrations of ethanol for 7 days (chronic treatment) (10 mM-2.5 M; mean +/- SEM; ***p<0.001; one-way ANOVA followed by Dunnett’s multiple comparison test). Each dot represents values obtained from one well (n= 3 independent experiments) normalized to the corresponding UT controls exposed to the assay media only. **(F)** Schematic of MNH assay workflow of ethanol exposure for 1 hour (acute treatment). **(G)** Quantification of MMP in 4-8-week-old DA neurons from control iPSCs (XM001) treated with increasing concentrations of ethanol for 1 hour (acute treatment; mean +/- SEM; *p<0.05, ***p< 0.001; one-way ANOVA followed by Dunnett’s multiple comparison test). Each dot represents values obtained from one well (n= 3 independent experiments) normalized to UT controls exposed to the assay media only. **(H-J)** Quantification of neuronal profiling including neurite count **(H)**, neurite length **(I)**, and branch points **(J)** in 4-8-week-old DA neurons from control iPSCs (XM001) treated with increasing concentrations of ethanol for 1 hour (acute treatment) (mean +/- SEM; *p<0.05, **p<0.01, ***p<0.001; one-way ANOVA followed by Dunnett’s multiple comparison test). Each dot represents values obtained from one well (n= 3 independent experiments) normalized to the corresponding UT controls exposed to assay media only. **(K)** Quantification of the MMP in 4-8-week-old DA neurons from hESCs (H1 line) treated with increasing concentrations of ethanol for 1 hour (acute treatment; mean +/- SEM; ***p<0.001; one-way ANOVA followed by Dunnett’s multiple comparison test). Each dot represents values obtained from one well (n= 3 independent experiments) normalized to the corresponding UT controls exposed to the assay media only. **(L-N)** Quantification of neuronal profiling including neurite count **(L)**, neurite length **(M)**, and branch points **(N)** in 4-8-week-old DA neurons from hESCs (H1 line) treated with increasing concentrations of ethanol for 1 hour (acute treatment; mean +/- SEM; *p<0.05, ***p<0.001; one-way ANOVA followed by Dunnett’s multiple comparison test). Each dot represents values obtained from one well (n= 3 independent experiments) normalized to the corresponding UT controls exposed to the assay media only.

We next investigated whether the observed changes in mitochondrial neuronal health could be recapitulated in DA neurons after acute exposure to ethanol. We plated 4-8-week-old DA neurons generated form iPSCs (XM001) and hESCs (H1) on the assay plate, kept them for 5 days and then exposed them to ethanol for 1 hour before the assay **(Fig. 3F)**. Acute ethanol exposure led to a dose-dependent reduction of MMP starting at a concentration of 500 mM ethanol in DA neurons derived from iPSCs **(Fig. 3G**) and hESCs **(Fig. 3K)**. Acute ethanol exposure caused a parallel dose-dependent reduction in neurite count and neurite length starting at 500 mM ethanol in DA neurons derived from iPSCs **(Fig. 3H-I)** and hESCs **(Fig. 3L-M)**, and a dose-dependent decrease in branch points starting at 500 mM ethanol in DA derived from iPSCs **(Fig. 3J)** and at 1 M ethanol in DA neurons derived from hESCs **(Fig. 3N)**.

Altogether, using the MNH assay to quantify ethanol-induced neurotoxicity, we determined that acute exposure to ethanol for 1 hour was sufficient to recapitulate the negative effects on mitochondrial neuronal health, which we observed in DA neurons after 7 days of chronic ethanol exposure. Neurotoxicity caused by ethanol ingestion may thus occur very rapidly. Therefore, in order to prevent ethanol-induced neurotoxicity, we might need to identify strategies to counteract the acute consequences of ethanol exposure.

### DA neurons from AUD patients recapitulate acute ethanol-induced neurotoxicity

Next, we aimed to address ethanol-induced neurotoxicity in DA neurons derived from individuals diagnosed with AUD according to DSM-5 and ICD-10 (alcohol dependence). Using Sendai viruses-mediated reprogramming, we generated iPSCs from four AUD patients: BIH232, BIH234, BIH235, and BIH236. All AUD-iPSCs expressed the pluripotency-associated protein markers OCT4 and TRA1-60 **(Fig. 4A)** and exhibited normal karyotypes **(Fig. 4C)**. Compared to somatic fibroblasts, AUD-iPSCs upregulated pluripotency-associated gene markers *OCT4, NANOG, SOX2, DNMT3B* and *DPPA4*, and downregulated the fibroblast-associated gene marker *VIM* **(Fig. 4B)**. AUD-iPSCs were pluripotent, as they were capable of generating cells belonging to the three germ layers: endoderm, mesoderm, and ectoderm **(Fig. 4D)**. We then differentiated AUD-iPCSs into NPCs and DA neurons. AUD-NPCs expressed NPC-specific protein markers NESTIN and PAX6 **(Fig. 5A)**. AUD-DA neurons expressed the pan-neuronal protein marker TUJ1 and the dopaminergic protein marker TH **(Fig. 5A)**.

**Fig. 4.**
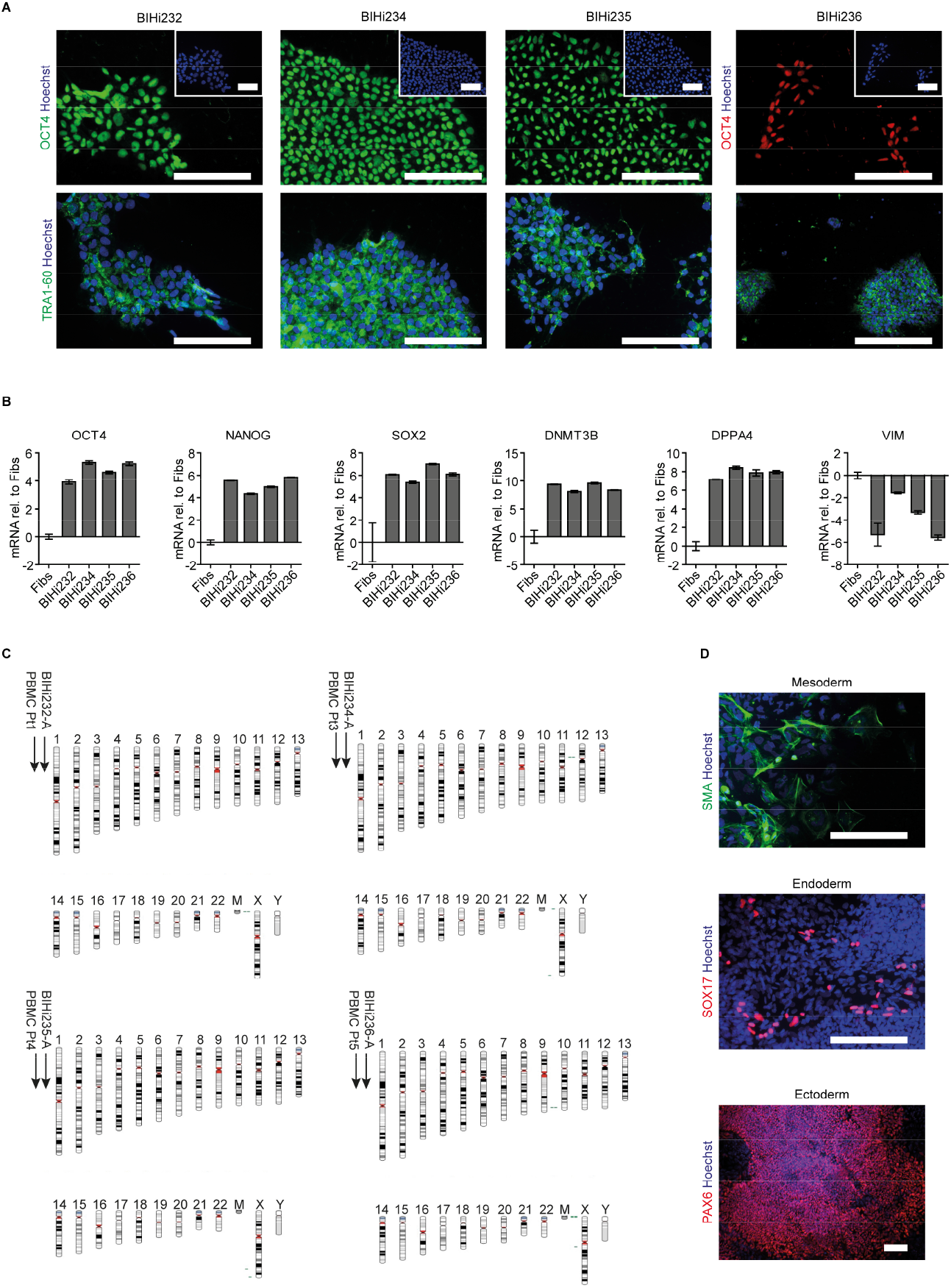
Characterization of AUD patient-derived iPSCs. **(A)** Representative immunostaining images of pluripotency-associated markers OCT4 and TRA1-60 in AUD-patient derived iPSCs (BIHi232, BIHi234, BIHi235, and BIHi236). We counterstained the cells using Hoechst. Scale bar: 100 μm. **(B)** Quantitative real-time RT-PCR analysis of pluripotency-associated markers in AUD patient-derived iPSCs (BIHi232, BIHi234, BIHi235, and BIHi236) relative to fibroblasts (Fibs; CON2). Relative transcript levels of each gene were calculated based on the 2-ΔΔCT method. Data were normalized to the housekeeping gene *GAPDH* and are presented as mean LOG2 ratios in relation to fibroblasts (mean +/- SD). **(C)** Single nucleotide polymorphism (SNP)-based virtual karyotype of AUD patient-derived iPSCs. We compared the karyotype of the iPSCs to the corresponding parental cells (peripheral blood mononuclear cells, PBMCs). We did not see any larger areas of insertions or deletions. Green: area with genomic gain; red: with genomic loss; gray: area with loss of heterozygosity. Pt= patient. **(D)** Representative immunostaining images of AUD-patient derived iPSCs differentiated into cells belonging to the three germ layers: mesoderm (smooth muscle actin, SMA), endoderm (SOX17), and ectoderm (PAX6). Scale bar: 100 μm.

**Fig. 5.**
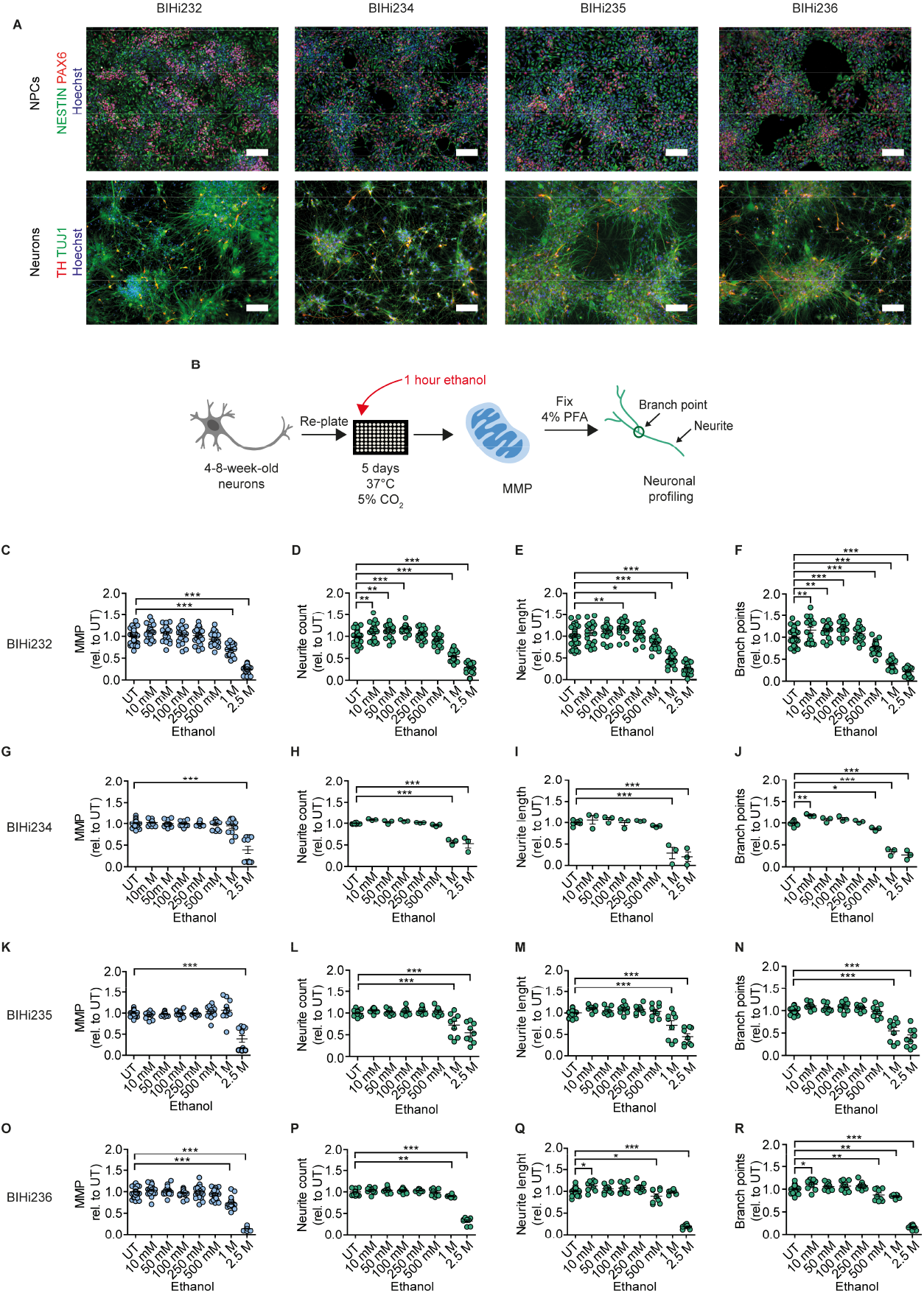
Acute ethanol exposure leads to decreased MMP and reduced branching complexity in AUD-patient iPSC-derived DA neurons. **(A)** Above: representative immunostaining images of NPC-associated markers NESTIN and PAX6 counterstained with Hoechst in NPCs from AUD-patient derived iPSCs (BIHi232, BIHi234, BIHi235, and BIHi236). Scale bar: 100 μm. Below: representative immunostaining images of neuron-associated marker TUJ1 and dopaminergic neuron-associated marker TH counterstained with Hoechst in DA neurons from AUD-patient derived iPSCs (BIHi232, BIHi234, BIHi235, and BIHi236). Scale bar: 100 μm. **(B)** Schematic of MNH assay workflow of ethanol exposure for 1 hour (acute treatment) on AUD patient iPSC-derived DA neurons. **(C-R)** MNH assay-based quantification of MMP **(C;G;K;O)** and branching complexity **(D-F; H-J; L-N; P-R)** in 4-8-week-old DA neurons from AUD patient-derived iPSCs (**C-F** BIHi232; **G-J** BIHi234; **K-N** BIHi235; **O-R** BIHi236) treated with increasing concentrations of ethanol for 1 hour (acute treatment; mean +/- SEM; *p<0.05, **p<0.01, ***p<0.001; one-way ANOVA followed by Dunnett’s multiple comparison test). Each dot represents values obtained from one well (n= 3 independent experiments) normalized to the corresponding UT controls.

We applied the MNH assay on AUD-DA neurons to quantify changes in MMP and neuronal arborization following 1 hour of acute ethanol exposure **(Fig 5B)**. Acute ethanol exposure decreased MMP in AUD-DA neurons as observed in healthy DA neurons, but only at high concentrations of 1 M and 2.5 M for lines BIH232 **(Fig. 5C)** and BIH236 **(Fig. 5O)**, and 2.5 M for lines BIH234 **(Fig. 5G)** and BIH235 **(Fig. 5K)**. Acute ethanol exposure also caused a dose-dependent reduction in neurite outgrowth that was variable among the AUD-DA neurons, starting at 1 M ethanol for neurite count **(Fig. 5D; 5H; 5L; 5P)**, 500 mM ethanol for neurite length **(Fig. 5E; 5Q)**, and 500 mM ethanol for branch points **(Fig. 5F; 5J; 5R)**.

Taken together, acute ethanol exposure negatively affected mitochondrial neuronal health in AUD-derived DA neurons. The individual AUD lines showed some level of heterogeneity, suggesting a different susceptibility to alcohol toxicity in different subjects. Nonetheless, the pattern of neurotoxicity detected by the MNH assay in AUD-DA neurons recapitulated the one observed in control DA neurons, suggesting that control neurons can be used to carry out studies aiming at identifying strategies to counteract ethanol-induced neurotoxicity.

### MNH-based proof-of-concept compound screening identified modulators of ethanol neurotoxicity

We next sought to determine whether the MNH assay could be used to carry out compound screenings to identify modulators of neurotoxicity. For screening, we used DA neurons derived from control iPSCs (XM001 line). We focused our attention on acute ethanol exposure. Using a library containing 700 FDA-approved drugs (Selleckchem # z65122), we selected 48 compounds that are approved for clinical use in the context of neurological diseases (compound numbers 1-48) **(Fig. 6B-D**; **Table 1**). Additionally, we included 5 drugs that are used for the treatment of AUD patients (compound numbers 49-53) **(Fig. 6B-D**; **Table 1**).

**Fig. 6.**
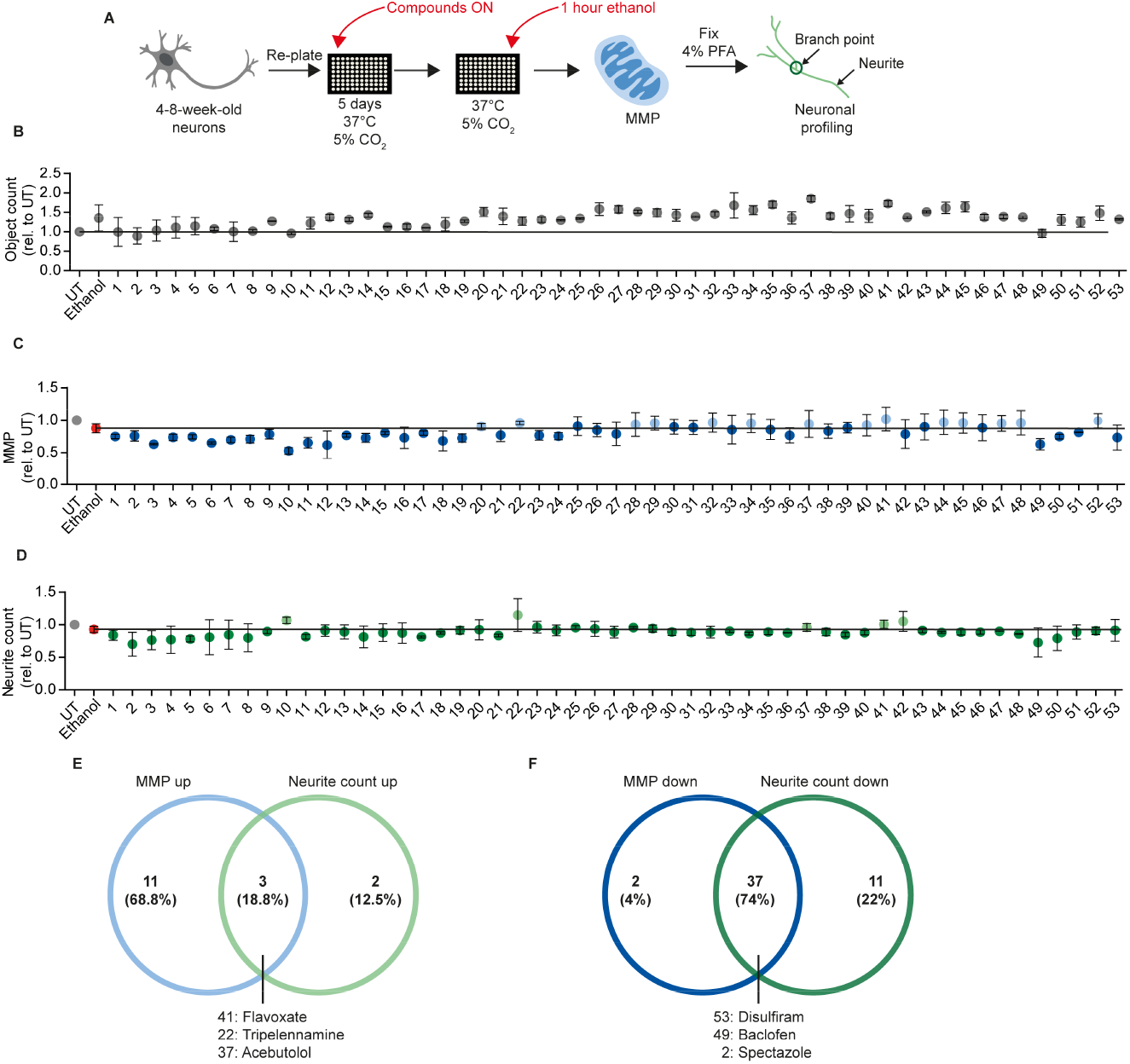
Proof-of-concept compound screening in human DA neurons exposed to ethanol. **(A)** Schematic MNH assay workflow for compound screening. **(B)** MNH assay-based quantification of object count of DA neurons from control iPSCs (XM001 line) after ON treatment with 48 FDA-approved drugs (Selleckchem, z65122) and 5 AUD drugs with subsequent exposure to 1 M ethanol for 1 hour (acute treatment; mean +/- SD; n= 2 independent experiments). Values from each experiment were normalized to the corresponding UT controls. The black line marks the mean of UT control. **(C-D)** MNH assay-based quantification of MMP **(C)** and neuronal profiling (**D**; neurite count) after ON treatment with 48 FDA-approved drugs out of the Selleckchem library (z65122) and 5 AUD drugs with subsequent exposure to 1 M ethanol for 1 hour (acute treatment). Values from each experiment were normalized to the corresponding UT controls (mean +/- SD; n= 2 independent experiments). The black line marks the mean value of ethanol treatment. Above: MMP relative to UT. The light blue dots mark the positive hit compounds that increased MMP in presence of 1 M ethanol compared to only ethanol treated (1 M, red dot). The dark blue dots mark the negative hit compounds. Below: neurite count relative to UT. The light green dots mark the positive hit compounds that increased neurite count in presence of 1 M ethanol compared to DA neurons exposed to ethanol only (1 M, red dot). The dark green dots mark the negative hit compounds. **(E)** Venn diagram for the compounds that increased MMP (“MMP up”) and increased neuronal outgrowth (“neurite count up”). **(F)** Venn diagram for the compounds that decreased the MMP (“MMP down”) and the neuronal outgrowth (“neurite count down”).

**Table 1.** List of selected compounds from the FDA-approved drug screening library (z65122)

| **compound**<br>**#** | **Name** | **M.w.** | **Target** | **Indication** | **Brief Description** | **CAS**<br>**Number** |
| --- | --- | --- | --- | --- | --- | --- |
| 1 | Bethanechol chloride | 196.68 | AChR | Neurological<br>Disease | Selective muscarinic receptor agonist without any effect on nicotinic receptors. | 590-63-6 |
| 2 | Econazole nitrate<br>(Spectazole) | 444.7 | Anti-infection,<br>Calcium<br>Channel | Neurological<br>Disease | Ca<sup>2+</sup> channel blocker, used as an antifungal medicine that fights infections caused by fungus. | 24169-02-6 |
| 3 | Acetanilide<br>(Antifebrin) | 135.16 | n.d | Neurological<br>Disease | Aniline derivative and has possess analgesic. | 103-84-4 |
| 4 | Cefprozil hydrate<br>(Cefzil) | 407.44 | n.d | Neurological<br>Disease | Second-generation cephalosporin type antibiotic. | 121123-17-9 |
| 5 | Tolterodine tartrate<br>(Detrol LA) | 475.57 | AChR | Neurological<br>Disease | Tartrate salt of tolterodine that is a competitive muscarinic receptor antagonist. | 124937-52-6 |
| 6 | Azelastine<br>hydrochloride<br>(Astelin) | 418.36 | Histamine<br>Receptor | Neurological<br>Disease | Potent, second-generation, selective,<br>histamine antagonist. | 79307-93-0 |
| 7 | 5-Aminolevulinic<br>acid hydrochloride | 167.59 | n.d | Neurological<br>Disease | Intermediate in heme biosynthesis in the<br>body and the universal precursor of<br>tetrapyrroles. | 5451-09-2 |
| 8 | Ronidazole | 200.15 | Anti-infection | Neurological<br>Disease | Antiprotozoal agent. | 7681-76-7 |
| 9 | Miglitol (Glyset) | 207.22 | Carbohydrate<br>Metabolism | Neurological<br>Disease | Oral anti-diabetic drug. | 72432-03-2 |
| 10 | Buflomedil HCl | 343.85 | Adrenergic<br>Receptor | Neurological<br>Disease | Vasodilator used to treat claudication or the<br>symptoms of peripheral arterial disease. | 35543-24-9 |
| 11 | PCI-32765<br>(Ibrutinib) | 440.5 | Src | Neurological<br>Disease | Potent and highly selective Btk inhibitor<br>with. | 936563-96-1 |
| 12 | Amoxicillin<br>(Amoxycillin) | 365.4 | Anti-infection | Neurological<br>Disease | Moderate-spectrum, bacteriolytic, β-lactam<br>antibiotic. | 26787-78-0 |
| 13 | Cabazitaxel (Jevtana) | 835.93 | Microtubule<br>Associated | Neurological<br>Disease | Semi-synthetic derivative of a natural taxoid. | 183133-96-2 |
| 14 | Niclosamide<br>(Niclocide) | 327.12 | STAT | Neurological<br>Disease | Inhibits DNA replication and inhibits STAT. | 50-65-7 |
| 15 | Lurasidone HCl | 529.14 | Dopamine<br>Receptor | Neurological<br>Disease | Antipsychotic, inhibits Dopamine D2, 5-HT2A, 5-HT7, 5-HT1A and noradrenaline α2C. | 367514-88-3 |
| 16 | D-Phenylalanine | 165.19 | n.d | Neurological<br>Disease | Carboxypeptidase A, endorphinase and enkephalinase inhibitor, enhances endorphin production and diminishes pain. | 673-06-3 |
| 17 | Palonosetron HCl | 332.87 | 5-HT Receptor | Neurological<br>Disease | 5-HT3 antagonist used in the prevention and treatment of chemotherapy-induced nausea and vomiting. | 135729-62-3 |
| 18 | Azelnidipine | 582.65 | Calcium<br>Channel | Neurological<br>Disease | Dihydropyridine calcium channel blocker. | 123524-52-7 |
| 19 | Dexmedetomidine | 200.28 | Adrenergic<br>Receptor | Neurological<br>Disease | Sedative medication used by intensive care units and anesthesiologists. | 113775-47-6 |
| 20 | Etravirine (TMC125) | 435.28 | Reverse<br>Transcriptase | Neurological<br>Disease | Non-nucleoside reverse transcriptase inhibitor (NNRTI) used for the treatment of HIV. | 269055-15-4 |
| 21 | Oxybutynin chloride | 393.95 | AChR | Neurological<br>Disease | Anticholinergic medication used to relieve urinary and bladder difficulties. | 1508-65-2 |
| 22 | Tripelennamine HCl | 291.82 | Histamine<br>Receptor | Neurological<br>Disease | H1 antagonist, inhibiting PhIP glucuronidation | 154-69-8 |
| 23 | Articaine HCl | 320.84 | n.d | Neurological<br>Disease | Dental local anesthetic which contains an additional ester group that is metabolized by esterases in blood and tissue. | 23964-57-0 |
| 24 | Doxylamine<br>Succinate | 388.46 | Histamine<br>Receptor | Neurological<br>Disease | Competitively inhibits histamine at H1 receptors with substantial sedative and anticholinergic effects. | 562-10-7 |
| 25 | Atomoxetine HCl | 291.82 | 5-HT Receptor | Neurological<br>Disease | Selective norepinephrine (NE) transporter inhibitor with $K_i$ of 5 nM, with 15- and 290-fold lower affinity for human 5-HT and DA transporters. | 82248-59-7 |
| 26 | Brinzolamide | 383.51 | Carbonic<br>Anhydrase | Neurological<br>Disease | Potent carbonic anhydrase II inhibitor. | 138890-62-7 |
| 27 | Ropinirole HCl | 296.84 | Dopamine<br>Receptor | Neurological<br>Disease | Selective dopamine D2 receptors inhibitor. | 91374-20-8 |
| 28 | Azlocillin sodium<br>salt | 484.48 | Anti-infection | Neurological<br>Disease | Acylampicillin with a broad spectrum against bacteria. | 37091-65-9 |
| 29 | Azacyclonol | 267.37 | 5-HT receptor | Neurological<br>Disease | Drug used to diminish hallucinations in psychotic individuals. | 115-46-8 |
| 30 | Reboxetine mesylate | 409.5 | n.d | Neurological<br>Disease | Norepinephrine reuptake inhibitor. | 98769-84-7 |
| 31 | Meptazinol HCl | 269.81 | Opioid Receptor | Neurological<br>Disease | Ppoid analgesic, which inhibits [$^3$H]dihydromorphine binding. | 59263-76-2 |
| 32 | Nalmefene HCl | 375.89 | Anti-infection | Neurological<br>Disease | Naltrexone analogue with opioid antagonistic property. | 58895-64-0 |
| 33 | Amidopyrine | 231.29 | n.d | Neurological<br>Disease | Analgesic, anti-inflammatory, and antipyretic properties. | 58-15-1 |
| 34 | Moclobemide | 268.74 | MAO | Neurological<br>Disease | MAO-A (5-HT) inhibitor. | 71320-77-9 |
| 35 | Lithocholic acid | 376.57 | FXR | Neurological<br>Disease | Acts as a detergent to solubilize fats for absorption. | 434-13-9 |
| 36 | Ethambutol HCl | 277.23 | Anti-infection | Neurological<br>Disease | Bacteriostatic antimycobacterial agent, which obstructs the formation of cell wall by inhibiting arabinosyl transferases. | 1070-11-7 |
| 37 | Acebutolol HCl | 372.89 | Adrenergic<br>Receptor | Neurological<br>Disease | β-adrenergic receptors antagonist used in the treatment of hypertension, angina pectoris and cardiac arrhythmias. | 34381-68-5 |
| 38 | Hyoscyamine<br>(Daturine) | 289.37 | AChR | Neurological<br>Disease | AChR inhibitor. | 101-31-5 |
| 39 | Procaine<br>(Novocaine) HCl | 272.77 | Sodium Channel | Neurological<br>Disease | Inhibitor of sodium channel, NMDA receptor and nAChR with IC<sub>50</sub> of 60 µM, 0.296 mM and 45.5 µM, which is also an inhibitor of 5-HT<sub>3</sub> with K<sub>D</sub> of 1.7 µM. | 51-05-8 |
| 40 | Dicyclomine HCl | 345.95 | M1 and M3<br>muscarinic<br>receptors<br>antagonist | Neurological<br>Disease | Anticholinergic tertiary amine. | 67-92-5 |
| 41 | Flavoxate HCl | 427.92 | AChR | Neurological<br>Disease | Muscarinic AChR antagonist. | 3717-88-2 |
| 42 | Aclidinium Bromide | 564.55 | AChR | Neurological<br>Disease | Human muscarinic AChR M1, M2, M3, M4 and M5. | 320345-99-1 |
| 43 | Bismuth<br>Subsalicylate | 362.09 | COX | Neurological<br>Disease | Active ingredient in Pepto-Bismol and inhibits prostaglandin G/H Synthase 1/2. | 14882-18-9 |
| 44 | Diphepanil<br>methanesulfate | 389.51 | AChR | Neurological<br>Disease | Quaternary ammonium anticholinergic, it binds muscarinic acetylcholine receptors (mAChR). | 62-97-5 |
| 45 | Doxapram HCl | 432.98 | TASK-1, TASK-3, TASK-1/TASK-3 | Neurological<br>Disease | TASK-1, TASK-3, TASK-1/TASK-3 heterodimeric channel function | 7081-53-0 |
| 46 | Methazolamide | 236.27 | Carbonic Anhydrase | Neurological<br>Disease | Carbonic anhydrase inhibitor. | 554-57-4 |
| 47 | Bextra (valdecoxib) | 314.36 | COX | Neurological<br>Disease | Potent and selective inhibitor of COX-2. | 181695-72-7 |
| 48 | Primidone<br>(Mysoline) | 218.25 | Sodium channel | Neurological<br>Disease | Anticonvulsant of the pyrimidinedione class. | 125-33-7 |
| 49 | Baclofen | 213,661 | GABA receptor | AUD drug | Derivative of the neurotransmitter γ-aminobutyric acid (GABA). | 1134-47-0 |
| 50 | Naltrexone<br>hydrochloride | 341.401 | Opioid receptor | AUD drug | N-cyclopropylmethyl derivative of<br>oxymorphone. Competitive antagonists at<br>the μ-opioid receptor (MOR). | 16590-41-3 |
| 51 | Acamprosate calcium | 181.211 | GABA receptor | AUD drug | NMDA receptor antagonist and positive<br>allosteric modulator of GABA<sub>A</sub> receptors. | 77337-76-9 |
| 52 | Clomethiazole<br>hydrochloride | 161.653 | GABA receptor | AUD drug | Positive allosteric modulator at the<br>barbiturate/picrotoxin site of the GABA<sub>A</sub><br>receptor. | 533-45-9 |
| 53 | Disulfiram | 296.52 | Dehydrogenase | AUD drug | Inhibitor of the enzyme acetaldehyde<br>dehydrogenase. | 97-77-8 |

We plated 4-8-week-old DA neurons and cultured them for 5 days. We treated DA neurons overnight (ON) with 0.5 μM of compound for each of the 53 compounds in a proof-of-concept experiment. We then exposed DA neurons to 1 M ethanol for 1 hour and performed the MNH assay **(Fig. 6A)**. Total object count, obtained by quantifying Hoechst-stained nuclei, showed no significant changes in cell number following the compound treatments, suggesting that we could confidently compare treated samples to untreated samples (**Fig. 6B**). As baseline for comparisons, we used DA neurons treated for 1 hour with 1 M ethanol that were not pre-exposed to any compound **(Fig. 6C-D, red dot)**. We defined a compound as a positive hit for MMP or neurite count when the compounds led to values that were higher than the baseline **(Fig. 6C-D, black horizontal line)**. We defined a compound as a negative hit for MMP or neurite count when the compounds led to values that were similar or lower than the baseline.

14 positive hit compounds led to higher MMP in the presence of ethanol **(Fig. 6C, light blue dots)**, and 5 positive hit compounds increased the number of neurites in the presence of ethanol **(Fig. 6D, light green dots)**. 3 positive hit compounds ameliorated both MMP and neurite count: flavoxate, tripelennamine, and acebutolol **(Fig. 6E)**. 39 negative hit compounds had either no effect or decreased MMP compared to ethanol **(Fig. 6C, dark blue dots)**, and 48 negative hit compounds had either no effect or decreased neurite count compared to 1 M ethanol **(Fig. 6D, dark green dots)**. 37 negative hit compounds decreased both MMP and neurite count in the presence of ethanol, suggesting enhanced neurotoxicity **(Fig. 6F)**. Among these 37 negative hit compounds, we identified spectazole, disulfiram, and baclofen **(Fig. 6F)**. Interestingly, these latter 2 drugs are commonly employed in AUD treatment.

We then aimed to assess whether the modulators of mitochondrial neuronal health identified in the screening could exert a modulatory effect on DA neurons independently of ethanol. We selected flavoxate and disulfiram as a positive hit compound and a negative hit compound, respectively. We used 4-8-week-old DA neurons from control iPSCs (XM001 line), re-plated them, and kept them for 5 days before ON treatment with either flavoxate or disulfiram **(Fig. 7A)**. DA neurons treated with 1 μM flavoxate showed a beneficial effect on MMP **(Fig. 7B)** and neurite arborization **(Fig. 7C-E)** compared to the DMSO-treated DA neurons. Conversely, disulfiram negatively affected the MMP **(Fig. 7F)** as well as the branching complexity **(Fig. 7G-I)** of DA neurons at concentrations between 0.5 and 10 μM. These data confirm the results of the screening and suggest that by identifying compounds counteracting neurotoxicity, it may be possible to discover general modulators of mitochondrial neuronal health.

**Fig. 7.**
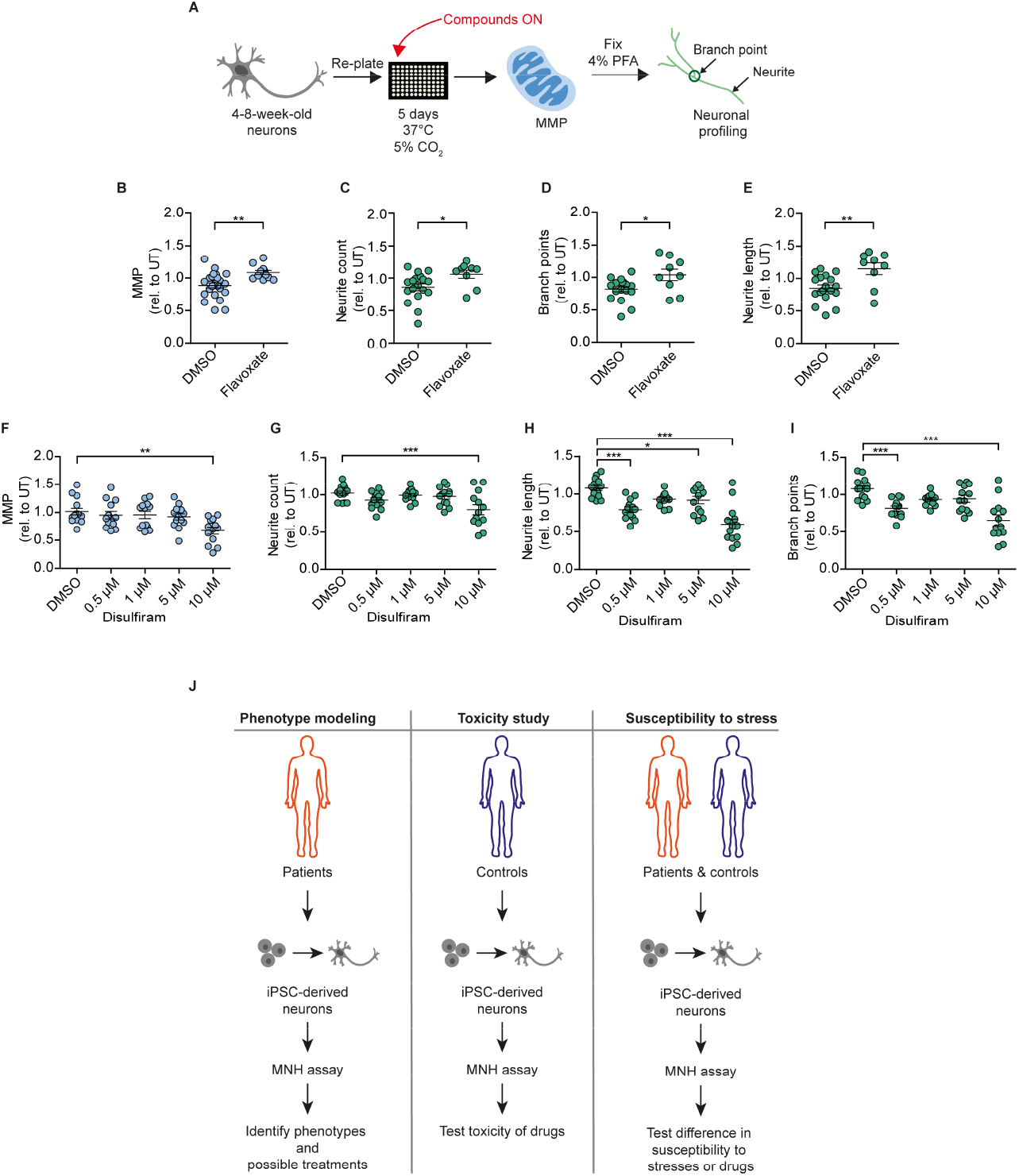
Compounds modulating neuronal toxicity in human DA neurons. **(A)** Schematic MNH assay workflow for neuronal toxicity. **(B-E)** MNH assay-based quantification of MMP (**B**) and neuronal profiling including neurite count (**C**), neurite length (**D**), and branch points (**E**) after ON treatment of 4-8-week-old DA neurons from control iPSCs (XM001) with 1 μM flavoxate (n= 3 independent experiments; mean +/- SEM; *p<0.05, **p<0.01; one-way ANOVA followed by Dunnett’s multiple comparison test). **(F-I)** MNH assay-based quantification of MMP (**F**) and neuronal profiling including neurite count (**G**), neurite length (**H**), and branch points (**I**) after ON treatment of 4-8-week-old DA neurons from control iPSCs (XM001) with increasing concentrations of disulfiram (n= 3 independent experiments; mean +/- SEM; *p<0.05, **p< 0.01; ***p<0.001; one-way ANOVA followed by Dunnett’s multiple comparison test). **(J)** Schematics of potential applications of the MNH assay.

## Discussion

In this study, we developed a method - the MNH assay - to assess the cellular health of human neurons derived from human pluripotent stem cells. We employed the MNH assay to dissect the neurotoxicity induced by ethanol on DA neurons. Previous iPSC-based studies assessed alcohol-mediated transcriptional changes in NMDA receptor expression (27), in GABA receptor expression (28), and in the expression of genes involved in cholesterol homeostasis, notch signaling, and cell cycle (29). Ethanol exposure for 1 day or 7 days was found to negatively influence the generation of mature neurons from NPCs (30). However, the effect of ethanol on mitochondrial neuronal function and neurite outgrowth remained to be determined. Using the MNH assay, we showed that acute exposure to ethanol was sufficient to cause loss of mitochondrial neuronal health, similarly to the one happening after 7 days of chronic ethanol exposure. Importantly, acute ethanol-induced mitochondrial and neuronal toxicity occurred in a comparable manner in different DA neurons derived from control individuals and from subjects diagnosed with AUD.

Using the MNH assay, we identified potential modulators of ethanol-induced neurotoxicity. Within a proof-of-concept screening, out of the 53 tested compounds flavoxate, tripelennamine, and acebutolol were found to potentially positively affect mitochondrial health and neurite outgrowth capacity. Conversely, 37 compounds showed a potential worsening effect in the presence of ethanol. Further studies should be carried out to confirm the exploratory results and determine the pharmacokinetic and pharmacodynamic characteristics of the hit candidates. Most drugs of abuse, including alcohol, have significant neurotoxic effects (31). Understanding the level of neurotoxicity caused by ethanol may enable the development of protective drugs to reduce the risk of developing neurodegenerative effects as consequence of excessive alcohol consumption. Interestingly, two drugs commonly employed in subjects with AUD, disulfiram and baclofen, negatively modulated mitochondrial neuronal health in the presence of ethanol. These findings warrant further exploration and raise concerns with respect to the potentially enhanced toxicity of drugs to treat AUD when used in concert with alcohol intake. Among the hits identified by the screening, we discovered possible modulators of mitochondrial neuronal health independent of ethanol. Flavoxate positively affected mitochondrial neuronal health, whereas disulfiram negatively affected mitochondrial neuronal health. In-depth analyses are warranted to confirm their function for human neuronal health. More generally, the findings suggest that the MNH assay might be used for various applications, including disease-specific studies, neurotoxicity studies, and studies aimed to assess the individual neuronal susceptibility to different stimuli (**Fig. 7J**).

The assessment of possible toxic effects of compounds is of high importance in the drug development process. Neurotoxicity testing commonly relies on *in vivo* animal studies that are expensive and may not fully recapitulate the toxicity profile of humans (32,33). Recent studies used MEA to investigate the effect of compounds on spontaneous neuronal activity and network activity (34). Various HT screening methods have also been established (35–41). Nonetheless, there is a need for additional *in vitro* systems centered on human iPSC derivatives in order to capture different aspects of human neurotoxicity (42). We suggest the MNH assay as a novel HT assay centered on human neurons that can be further multiplexed with additional imaging-based techniques, including for example the addition of an apoptotic dye to quantify cell death (35,36). iPSC-derived 3D neural cultures may also be employed to assess the neurotoxic potential of drugs and environmental toxicants (43). Therefore, future implementations should aim to adapt the MNH assay to 3D culture systems, such as iPSC-derived brain organoids (44).

Taken together, we developed a novel assay that enabled us to capture the ethanol-induced toxicity in human DA neurons and to identify potential small molecule modulators. We propose the MNH assay as a tool for the evaluation of human neuronal health and for conducting HT drug discovery and drug toxicity studies.

## Author contributions

Conceptualization: J.P., A.P. Methodology: A.P., J.P., A.Z. Investigation: A.Z., J.C., N.S.T, S.D. Resources: A.P., J.P., E.W. Writing – Original Draft: A.Z., A.P., J.P. Writing – Review and Editing: A.Z., A.P., J.P., A.H. Supervision: A.P., J.P. Visualization: A.Z., A.P. Funding Acquisition: A.P., J.P., A.H.

## Acknowledgements

We thank members of the Prigione group for help and support with the cultures of iPSCs and derived neurons. We also thank Laura Deadelow and Eike Jakob Sputh for help with the generation of AUD-iPSCs. We acknowledge support from the MDC (BOOST award to A.P.), the Deutsche Forschungsgemeinschaft (DFG) (PR1527/1-1 to A.P.; SFB/TRR 265 B04 to J.P.), the Berlin Institute of Health (BIH) (BIH Validation Funds to A.P.), and the German Federal Ministry of Education and Research (BMBF) (e:Bio young investigator grant AZ.031A318 and AZ.031L0211 to A.P.; AERIAL P1 to A.H. and J.P.), and the UK DRI (Momentum Award and Programme Award to J.P.).

## Methods

### Generation of iPSCs from PBMCs

This study was approved by the ethics committee of the Charité – Universitätsmedizin Berlin (EA1/206/15). Patient peripheral blood mononuclear cells (PBMCs) were obtained after informed consent. In brief, blood collections were performed using standard, 8 ml Vacutainer Cell Processing Tubes (both sodium citrate and sodium heparin-based tubes are acceptable; BD Biosciences; Franklin Lakes, NJ USA). Samples were processed within 24 hours after blood collection by collecting the PBMC-containing upper phase and washed with ice-cold PBS. Cells were either frozen down or used directly for enrichment of the erythroblast population. In brief, cell pellet of 1 x10^6^ – 2 x10^6^ PBMCs was resuspended in 2 ml erythroblast cell expansion media containing basal blood media Stempo34 SFM (Thermo Fisher) supplemented with 1x Stempo34 nutrient supplement (Thermo Fisher), 1x Glutamine (Thermo Fisher) and cytokines such as 20 ng/ml IL3 (PeproTech), 20 ng/ml IL6 (PeproTech), 100 ng/ml Flt3 (PeproTech), 100 ng/ml SCF (PeproTech), 2 U/ml erythropoietin (EPO, Millipore) and plated in a standard 12-well plate (2 mL/well). The media was changed every other day and after 4 to 6 days a significant enrichment of highly proliferating erythroblast population was observed. This highly proliferative cell population was used for the reprogramming. Erythroblast cells from PBMCs were reprogrammed using the Cytotune^®^-iPS 2.0 Sendai Reprogramming Kit according to the manufacturer’s instructions. In brief, cells were transduced with CytoTune 2.0 Sendai virus coding for *OCT3/4, KLF4, SOX2* and *MYC* genes with recommended MOI in 1 ml of erythroblast cell expansion media containing 10 μg/ml Polybrene (Sigma). After 24 hours of transduction the virus was removed, and cells were cultured for further two days in erythroblast cell expansion media. From day 1 to 3 the cells were cultured in repro media 1 consisting of erythroblast cell expansion media, 200 μM/L sodium butyrate (NaB, Stem Cell Technologies) and 50 μg/ml ascorbic acid (Sigma). On day 3 after transduction 1 x10^5^ cells were plated onto mouse embryonic fibroblast (MEFs) layer or Vitronectin coated plate in repro media 1. From day 4 until day 7, the media was changed with repro media 2 containing basal blood media, NaB and ascorbic acid without cytokines. On day 7 the media was switched to repro media2 and E7 reprogramming media (1: 1) containing E6 basal media (Thermo Fisher) with NaB, ascorbic acid and 100 ng/ml FGF2 (PeproTech) and the same was used every day until day 13. From day 13 onwards the cells were cultured in E8 life (Thermo Fisher) until day 20. Emerging colonies were picked three weeks after the transduction and tested for remaining viral RNA expression using RT-PCR by checking the expression of Sendai viral reprogramming particles using primers specific to Sendai virus.

### iPSC culture

Control iPSCs were previously generated using episomal plasmids and described as XM001 (17). hESC line H1 was purchased from WiCell and was used in accordance with the German license of Alessandro Prigione issued by the Robert Koch Institute (AZ: 3.04.02/0077-E01). All PSCs were cultivated on Matrigel (BD Bioscience)-coated plates using: StemMACS iPS-Brew XF medium (Miltenyi Biotec GmbH, #130-104-368), supplemented with 0.1 mg/ml Pen/Strep (Thermo Fisher Scientific) and MycoZap [1x] (Lonza). We routinely monitored against mycoplasma contamination using PCR. 10 μM ROCK inhibitor (Enzo, ALX-270-333-M005) was added after splitting to promote survival. PSC cultures were kept in a humidified atmosphere of 5% CO_2_ at 37°C and 5% oxygen. All other cultures were kept under atmospheric oxygen condition. Pluripotency of the generated lines was confirmed following previously published procedures (45) using *in vitro* embryoid bodies (EB)-based differentiation. We reprogrammed patient PBMCs using Sendai viruses.

### SNP karyotyping

Briefly, genomic DNA was isolated using the DNeasy blood and tissue kit (Qiagen, Valencia, CA) and samples were analyzed using the human Illumina OMNI-EXPRESS-8v1.6 BeadChip. First, the genotyping was analyzed using GenomeStudio 1 genotyping module (Illumina). Thereafter, KaryoStudio 1.3 (Illumina) was used to perform automatic normalization and to identify genomic aberrations utilizing settings to generate B-allele frequency and smoothened Log R ratio plots for detected regions. The stringency parameters used to detect copy number variations (CNVs) were set to 75 kb (loss), 100 kb (gain) and CN-LOH (loss of heterozygosity) regions larger than 3 MB.

### Differentiation of NPCs and DA neurons

We obtained NPCs and DA neurons using a previously published protocol (19). Briefly, PSCs were detached from Matrigel-coated plates using StemPro^®^ Accutase^®^ Cell Dissociation Reagent (accutase; Thermo Fisher) and the collected sedimented cells were transferred into low-attachment petri dishes and kept for two days in ES-based medium containing: KO-DMEM [1x] (Gibco), KO-SR [1x] (Gibco), NEAA [1x] (Gibco), 2 mM L-Glutamine (Gibco), 0.1 mg/ml Pen/Strep (Gibco), 1 mM Pyruvate (Gibco) and MycoZap [1x] (Lonza), without FGF2, supplemented with 0.5 μM purmorphamine (PMA, Millipore), 3 μM CHIR 99021 (CHIR, Caymen Chemical), 10 μM SB-431542 (MACS Miltenyi) and 1 μm dorsomorphin (Sigma). From day 2 to day 4, the media was switched to: Neurobasal:DMEM/F12 [1:1], N2 [1x], B27 without vitamin A [1x], 2 mM L-Glutamine, 0.1 mg/ml Pen/Strep, 1 mM Pyruvate and MycoZap [1x] with the addition of 0.5 μM PMA, 3 μM CHIR 99021, 10 μM SB-431542 and 1 μM dorsomorphin. On day 4 the media was switched to the final maturation media containing: Neurobasal:DMEM/F12 [1:1], N2 [1x], B27 without vitamin A [1x], 2 mM L-Glutamine, 0.1 mg/ml Pen/Strep, 1 mM Pyruvate and MycoZap [1x], supplemented with 3 μM CHIR 99021, 0.5 μM PMA and 150 μM vitamin C (Sigma Aldrich). On day 6, the suspended cells were transferred onto Matrigel-coated well plates and kept in the maturation media with media exchange every 2 days. NPCs were maintained on this media without ROCK inhibitor and used for experiments between passage 7 and 20. For neuronal differentiation, we used NPCs between passage 7 and 13. To initiate the differentiation, the media was changed to: Neurobasal:DMEM/F12 [1:1], N2 [1x], B27 [1x], 2 mM L-Glutamine, 0.1 mg/ml Pen/Strep, 1 mM Pyruvate and MycoZap [1x] with addition of 200 μM vitamin C, 100 ng/mL FGF8 (R&D Systems) and 1 μM PMA. After 8 days, the media condition was replaced with: Neurobasal:DMEM/F12 [1:1], N2 [1x], B27 [1x], 2 mM L-Glutamine, 0.1 mg/ml Pen/Strep, 1 mM Pyruvate and MycoZap [1x] with the addition of 200 μM vitamin C, 0.5 μM PMA, 500 μM cAMP (StemCells), 10 ng/mL BDNF (MACS Miltenyi), 10 ng/mL GDNF (MACS Miltenyi) and 1 ng/mL TGFbeta3 (MACS Miltenyi). On day 9, cells were split with accutase and seeded on Matrigel-coated plates in: Neurobasal:DMEM/F12 [1:1], N2 [1x], B27 [1x], 2 mM L-Glutamine, 0.1 mg/ml Pen/Strep, 1 mM Pyruvate and MycoZap [1x] with the addition of 200 μM vitamin C, 500 μM cAMP, 10 ng/mL BDNF, 10 ng/mL GDNF, and 1 ng/mL TGFbeta3. The media was changed every 3-4 days and the differentiated cells were kept in culture for 4 weeks up to 8 weeks. 10 μM ROCK inhibitor (Enzo, ALX-270-333-M005) was always added after splitting to promote survival.

### PCR analyses

Gene expression analysis was performed by quantitative real-time RT-PCR (qRT-PCR) using SYBR Green PCR Master Mix and the ViiA™ 7 Real-Time PCR System (Applied Biosystems). For each target gene, cDNA samples and negative controls were measured in triplicates using 384-Well Optical Reaction Plates (Applied Biosystems). Relative transcript levels of each gene were calculated based on the 2−ΔΔCT method. Data were normalized to the housekeeping gene *GAPDH* and are presented as mean LOG2 ratios in relation to control cell lines. Primers were for OCT4 F: GTGGAGGAAGCTGACAACAA and R: ATTCTCCAGGTTGCCTCTCA, for NANOG F: CCTGTGATTTGTGGGCCTG and R: GACAGTCTCCGTGTGAGGCAT, for SOX2 F: GTATCAGGAGTTGTCAAGGCAGAG and R: TCCTAGTCTTAAAGAGGCAGCAAAC, for DNMT3B F: GCTCACAGGGCCCGATACTT and R: GCAGTCCTGCAGCTCGAGTTTA, for DPP4 F: TGGTGTCAGGTGGTGTGTGG and R: CCAGGCTTGACCAGCATGAA, for VIM F: GGAGCTGCAGGAGCTGAATG and R: GACTTGCCTTGGCCCTTGAG.

### Immunostaining

Cells grown on Matrigel-coated coverslips were fixed with 4% paraformaldehyde (PFA, Science Services) for 15 min at room temperature (RT) and washed two times with PBS. For permeabilization, cells were incubated with blocking solution containing 10% normal donkey serum (DNS) and 1% Triton X-100 (Sigma-Aldrich) in PBS with 0.05% Tween 20 (Sigma-Aldrich) (PBS-T) for 1 hour at RT. Primary antibodies were diluted in blocking solution and incubated overnight at 4°C on a shaker. Primary antibodies were used as follows: PAX6 (Covance, 1:200), SOX2 (Santa Cruz, 1:100), TUJ1 (Sigma-Aldrich, 1:3000), OCT4 (Santa Cruz, 1:300), TRA-1-60 (Millipore, 1:200), MAP2 (Synaptic System, 1:100), NANOG (R&D Systems, 1:200), SMA (DakoCytomation, 1:200), SOX17 (R&D Systems, 1:50), TH (Millipore, 1:300), FOXA2 (Sevenhills, 1:100). Corresponding secondary antibodies (Alexa Fluor, 1:300, Life Technologies) were diluted in blocking solution for 1 hour at RT on a shaker. Counterstaining of nuclei was carried out using 1:2,500 Invitrogen™ Hoechst 33342 (Hoechst; ThermoFisher). All images were acquired using the confocal microscope LSM510 Meta (Zeiss) in combination with the AxioVision V4.6.3.0 software (Zeiss) and further processed with AxioVision software and Photoshop CS6 version 6.2 (Adobe).

### Micro-electrode array (MEA) recordings

MEA recordings were conducted using the Maestro system from Axion BioSystems. A 48-well MEA plate with 16 electrodes per well was precoated with 1 mg/ml polyethyleminine (PEI; Sigma Aldrich) for 1 hour at RT, washed three times with sterile H20 and air-dried ON in the cell culture hood. Afterwards, the plate was pre-coated with Matrigel by placing a 7 μl drop of Matrigel (200 μg/ml) into the centre of the well (on top of the electrodes) and incubated for 1 hour at 37°C, 5% CO_2_. Remaining coating solution was removed, and 80,000 DA neurons per well were seeded on the electrodes in each well of the MEA plate by preparing a cell suspension of 7 μl and applying a droplet directly into the centre of each well of the MEA plate. Following an incubation of 1 hour at 37°C, 5% CO_2_ to ensure settlement of the DA neurons, 300 μl maturation media (for media formulation see “Differentiation of NPCs and DA neurons”) was slowly added to each well. Half of the media was refreshed every two days. Two weeks after plating, spontaneous activity was recorded at a sampling rate of 12.5 kHz under controlled conditions (37°C and 5% CO_2_) for 10 minutes on different days. Using Axion Integrated Studio (AxIS 1.4.1.9), a digital low pass filter of 4 kHz and high pass filter of 200 Hz cut-off frequency and a threshold of 5.5x the standard deviation was set to minimize false-positives detections. Raw data was analysed using the Neural Metric Tool (Axion BioSystems) to analyse spikes. Electrodes that measured more than 5 spikes per minute were considered active.

### High-content imaging (HCI)

HCI-based quantification of MMP and neuronal profiling was assessed using the HCI platform CellInsight CX7 microscope (Thermo Fisher Scientific) and the integrated BioApplications. Briefly, 4-8-week-old DA neurons were split using accutase and seeded at a density of 10,000-20,000 cells/well on Matrigel-coated 96 well plates with black-wall and clear-bottom (μClear, Greiner). The cells were live-stained with nonquenching concentrations of 0.5 nM TMRM (ThermoFisher) for 30 minutes at 37°C and 5% CO_2_ and counterstained with 1:10,000 Hoechst (Thermo Fisher). Parameters were set as follows: primary object detection (cell nuclei) was based on Hoechst staining, captured in channel 1. Detection of the cellular region of interest, captured in channel 2, was based on TMRM staining which accumulates in active mitochondria as a result of the MMP. Changes in TMRM intensity within the region of interest (ring region with adjusted threshold) were quantified using the “Cellomics CellHealthProfiling v4 BioApplication” (CellInsight CX7, Thermo Fisher Scientific) using the feature “MeanTargetAvgIntensityCh2”. Afterwards, neuronal cultures were fixed with 4% PFA for 15 minutes at RT, stained with TUJ1 antibody and counterstained with Hoechst (see “Immunostaining” for details on staining method). Parameters were set as follows: primary object detection (cell nuclei) was based on Hoechst staining, captured in channel 1. Detection of the region of interest (neurites) was based on TUJ1 staining, captured in channel 2. The morphological changes of TUJ1-postive cells were quantified using the “Cellomics Neuronal Profiling v4 BioApplication” (CellInsight CX7, Thermo Fisher Scientific). Features obtained with the BioApplication used for evaluation of the neuronal branching complexity were: “NeuriteTotalCountPerNeuronCh2”, “NeuriteTotalLengthPerNeuronCh2”, and “BranchPointTotalCountPerNeuronCh2”.

### Treatment of DA neurons

FCCP and Ant.A, or Oligomycin were applied to 4-8-week-old DA neurons together with the staining solution (0.5 nM TMRM and 1:10,000 Hoechst) and incubated for 30 minutes at 37°C, 5% CO_2_. Staining solution containing FCCP and Ant.A, or Oligomycin was removed and DA neurons were washed three times gently with 1x PBS before corresponding phenolred-free media was applied and cells were measured at the CellInsight CX7. Treatments with ethanol were performed 5 days after re-plating 4-8-week-old DA neurons on black-wall, round bottom 96-well plates (μClear, Greiner). 99.5% ethanol (v/v; Roth) was prediluted in culture media as 5 M stock solution freshly before use. Accordingly, 10 mM-2.5 M ethanol was further diluted in phenolred-free culture media and administered to the DA neurons. For the chronic ethanol exposure, freshly prepared culture media containing ethanol was changed every other day at the same time for 7 days including one day of withdrawal. Evaporation of ethanol following daily media changes in unsealed culture plates may provide a pattern of exposure more similar to alcohol consumption observed in human daily heavy drinkers. After 7 days of ethanol exposure, DA neurons were stained with TMRM and Hoechst (see details above). Following the removal of the staining solution, phenolred-free media containing ethanol was reapplied and incubated again for 1 hour before assessment of the MNH at the CellInsight CX7. For the acute ethanol exposure, DA neurons were first stained with TMRM and Hoechst (see details above) before freshly prepared phenolred-free culture media containing ethanol was administered to the DA neurons for 1 hour at 37°C, 5% CO_2_. After the incubation time, DA neurons where immediately processed at the CellInsight CX7. Drug treatment (flavoxate and disulfiram) was performed as described before on 4-8-week-old DA neurons with incubation ON at 37°C and 5% CO_2_. The next day, staining solution containing 0.5 nM TMRM and 1:10,000 Hoechst was added to the media for 30 min at 37°C, 5% CO_2_. Further processing was performed according to the MNH assay as described before.

### Compound screening – MNH assay

For the proof-of-concept compound screening, 48 compounds were selected from a library of 700 FDA-approved drugs (Selleckchem-z65122) and 5 drugs were added, used in the context of AUD treatment (**Table 1**). Briefly, 10,000-20,000 DA neurons/well were plated on black-wall, round bottom 96-well plates (μClear, Greiner) 5 days before adding the compounds ON. Compounds were used in a final concentration of 0.5 μM diluted in culture media. DMSO concentration was diluted down below 0.05%. The second day, DA neurons were stained with 0.5 nM TMRM together with 1:10,000 Hoechst diluted in phenol red-free culture medium for 30 min at 37°C, 5% CO_2_ by adding the staining solution into the media containing the drugs. After removal of the staining solution, compounds together with 1 M ethanol were applied to the DA neurons and incubated for 1 hour at 37°C, 5% CO_2_. Control wells included DA neurons kept untreated and DA neurons treated with 1 M ethanol only. HCI was conducted with the CellInsight CX7 microscope (Thermo Fisher) and analysed according to the “CellHealth Profiling” and “Neuronal Profiling” BioApplication. The same screening was repeated twice, and the values shown in **Fig. 6B-D** represent the mean values of the two runs (mean, +/- SD).

### Statistical analysis

Data were analyzed using GraphPad-Prism software (Prism 4.0, GraphPad Software, Inc.). Data presentation and respective statistical analysis of each individual graph are described in the respective figure legends. The z-factor is defined as the means (μ) and standard deviations (σ) of both the positive (p) and negative (n) controls as follows:

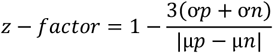

